# An immersive first-person navigation task for abstract knowledge acquisition

**DOI:** 10.1101/2020.07.17.208900

**Authors:** Doerte Kuhrt, Natalie R. St. John, Jacob L. S. Bellmund, Raphael Kaplan, Christian F. Doeller

## Abstract

Advances in virtual reality (VR) technology have greatly benefited spatial navigation research. By presenting space in a controlled manner, changing aspects of the environment one at a time or manipulating the gain from different sensory inputs, the mechanisms underlying spatial behaviour can be investigated. In parallel, a growing body of evidence suggests that the processes involved in spatial navigation extend to non-spatial domains. Here, we leverage VR technology advances to test whether participants can navigate abstract knowledge. We designed a two-dimensional quantity space - presented using a head-mounted display - to test if participants can navigate abstract knowledge using a first-person perspective navigation paradigm. To investigate the effect of physical movement, we divided participants into two groups: one walking and rotating on a motion platform, the other group using a gamepad to move through the abstract space. We found that both groups learned to navigate using a first-person perspective and formed accurate representations of the abstract space. Interestingly, navigation in the quantity space resembled behavioural patterns observed in navigation studies using environments with natural visuospatial cues. Notably, both groups demonstrated similar patterns of learning. Taken together, these results imply that both self-movement and remote exploration can be used to learn the relational mapping between abstract stimuli.

## Introduction

Virtual reality (VR) technology has made great advances in recent years. The use of life-like graphics, head-mounted displays (HMD) as well as omnidirectional motion platforms helps to create interactive environments in which participants behave naturally and intuitively. Researchers on the other hand, remain in full control and can track participants’ behaviour in great detail with advanced sensors. Consequently, VR has increasingly been used to study spatial navigation^1^, social interaction^2^ and multisensory integration^3^. It has also been used in therapeutic settings such as stroke rehabilitation^4^, pain therapy^5^ or phobia treatment^6^. In all of these fields, researchers make use of the interactive nature of VR paradigms: participants are fully immersed in a highly-controlled representation and interact with their environment in a safe yet realistic manner. They act from a first-person perspective, which creates a sense of presence and makes the environment relevant and stimulating.

Interactiveness, as facilitated by using VR, is also the central research topic in embodied learning. Research suggests that learning, even of abstract concepts or ideas, is aided by physically exploring them^7–9^. This has not only been shown in children, but also in adults^10–12^ and in a range of different tasks such as counting^13, 14^, language^15^, letter recognition^16, 17^, music concepts^18^ and understanding of physics^19^. Interacting with a concept increases its relevance, and makes it literally “graspable”. Interestingly, the effects of embodied learning have been shown on different levels of cognition that include passive viewing of images^20^, recognition memory^21^ and understanding of concepts^18, 19, 22^. The extent to which the way of learning shapes knowledge representation is still debated and may depend on the task at hand, but there are examples suggesting that the mode of acquisition has a lasting influence on knowledge representation^23, 24^. For example, even in educated adults, who no longer use their fingers for counting, number representation is linked to their finger counting technique during early childhood^14^. Physically representing knowledge in an interactive manner could therefore serve as a transformative pedagogical approach that extends beyond initial learning to influence how knowledge is maintained and retrieved.

One skill we naturally learn in an embodied fashion is spatial navigation. During navigation, we use multiple senses such as vision, vestibular and motor feedback to find our way. However, the necessity of individual senses during navigation remains debated, with several studies showing no differences in behaviour with varying amount of inputs^25–27^ while others show an effect on behavioural performance^28–33^. The exact reason for the discrepancy in these studies is unclear, but the field generally agrees that inputs from different senses are integrated^34–36^. Especially in ambiguous situations, multi-sensory integration reduces error and thus is particularly important when we are trying to navigate difficult terrain while keeping track of our point of origin and destination.

This idea of keeping track of where we are, where we came from and where we are going entails the formation of a representation of the environment, the so-called cognitive map as it was first introduced by Tolman^37^. The cognitive map can be used to flexibly guide behaviour to find known paths as well as unknown short-cuts. 70 years of investigating Tolman’s claims has led to the identification of several core mechanisms for spatial navigation that span across species (for review^38, 39^).

Beyond classical spatial navigation theories, a growing body of research suggests spatial navigation mechanisms are involved in processing more than physical space^40–44^. This entails the formation of a cognitive map of abstract domains organized in n-dimensions^41^. For example, imagine a task for which you have to judge the similarity between animals. Given a giraffe, a rabbit and a seagull, depending on which dimensions you consider, you will probably reach different conclusions about their similarity: seagulls and rabbits are more similar when you consider size of the animal, population number and habitat range, while rabbits and giraffes are more similar when you consider diet and taxonomy. Flexibly accessing knowledge and making decisions about similarity can thus be seen as estimating distances between points (animals) in a high-dimensional space (animal features), similar to how you would judge the distance between the town hall and the train station when you want to figure out how much time you will need to walk between these places.

Evidence for abstract cognitive spaces have been observed in social^45, 46^, semantic^47^, frequency^48^, odor^49^, abstract^50, 51^ and musical^52^ domains. These studies had participants navigate in the space and form associations with selected positions. Critically, all studies used the abstract domains only and did not use an intuitive visuospatial representation of the domain. The tasks used in these studies resemble physical space navigation in their dimensionality, but are usually missing a first-person perspective. During navigation of these abstract spaces, single-cell recordings have demonstrated the recruitment of spatially tuned cells^48, 49^. Consistently, modulations of BOLD-responses by inter-stimulus distances and grid-like hexadirectional signals have been observed in human fMRI^50, 51^.

What many of these studies are lacking is the first-person perspective resembling physical space navigation and the interactiveness by which we typically explore our environment. When navigating physical space, it is proposed that egocentric representations gained through a first-person perspective may be transformed to an allocentric reference frame to support a viewpoint independent map-like representation^53^. By creating an immersive abstract space that can be navigated from a first-person perspective, we can measure similar transformations in an abstract environment. This would then afford a greater comparison to the processing principles underlying physical navigation.

By combining findings from embodied learning with well-established behavioural patterns as present in spatial navigation, we aim to explore the idea that abstract knowledge can be represented in a map-like fashion and that it can be navigated using physical movement. Such a representational format would eliminate the hurdle of an intuitive spatial representation, since it could be applied to any type of knowledge that can be organized along a number of dimensions. More concretely, we want to build on the idea that physical movement, via multi-sensory integration, limits ambiguity and error in a difficult environment, which could be beneficial in abstract space navigation. Further, physical exploration of a concept has been shown to improve performance in embodied learning research^19^. We propose that a physically navigable conceptual space should therefore help participants build up a mental representation, which in turn should lead to better distance and direction estimations.

To test the idea, we created a two-dimensional abstract quantity space (figure 1 and supplementary video). This means participants saw quantities of 2 different geometric shapes: circles and rectangles. Each shape quantity defined the position along the respective axis of the space, so that each set of quantities encoded a position. Participants could manipulate the quantities to move between positions. To move in different directions, participants learned to connect a color-code to their facing direction in the space. We trained participants extensively to use the color and the quantities to navigate through this abstract space, and in the end, tested their ability to estimate distances and angles between positions, an ability supported by map-like representations of space. However, participants were never told that the abstract knowledge they were acquiring was spatialized by our task design. We deliberately refrained from explaining the relation between position and quantities as well as color and direction and avoided spatial instructions as much as possible throughout the experiment. Critically, we were interested in two main questions: can participants learn to navigate such an abstract space using color as directional and quantity as positional information? Second, are participants who navigate this space by physically exploring it more accurate in their distance and angle estimations than those who use a gamepad? Participants were randomly assigned to one of two groups. The movement group (MG) navigated the conceptual space by walking and rotating on an omni-directional motion platform while visual feedback was provided in a head-mounted display. The gamepad group (GG) viewed the space also using the head-mounted display and standing up, but the motion sensors were disabled, so that participants used a gamepad for rotations and translations in the space. Both groups underwent extensive behavioural testing, with several different measures to identify similarities and differences to physical space navigation.

**Figure 1.**
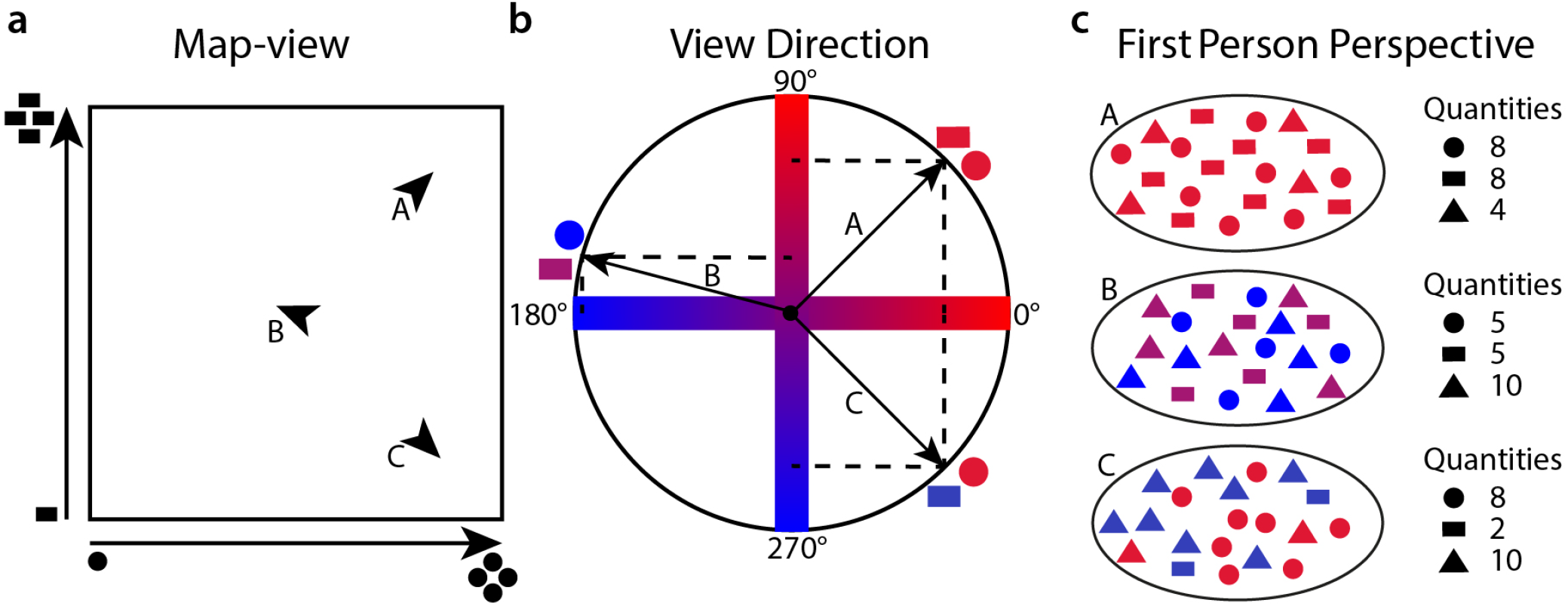
Quantity space. **a** Map-view representation of the two-dimensional space. The X-dimension is the quantity of circles, and the Y-dimension the quantity of rectangles presented to the participant. When a participant moves forward, the quantities of the items change. Arrows A, B and C represent example positions, with the arrow direction showing view direction. **b** We used color to encode view direction. The color-code ran between blue and red, with blue indicating a decrease, red indicating an increase and purple indicating no change in quantity if a participant were to move forward. For the X-dimension (circles) the cosine of the view direction determined the color, and for the Y-dimension, the sine of the view direction determined the color. While turning, the colors are continuously updated while the quantities stay stable. Color code for the view directions of positions A, B and C from **a** are shown as examples. **c** Down-sampled (5 %) representation of the first-person perspective when standing at the locations A, B and C, as presented in **a** and **b**. The participants only saw the space from a first-person perspective, and had to navigate using the quantities of items for positional information and colors for directional information. The triangles are filler items, so that each position always contained 400 items (in this example 20 items). Thus, visual complexity for the different positions was kept more stable. Triangles are dimension-specific: they share the color of the dimension they are “filling-up”. Numbers indicate the quantities of the items.

## Methods

### Participants

We used G*Power (3.1.9.2) to calculate the sample size per group aiming to achieve a power of 0.95 when using one-sided t-tests, with an assumed medium effect size of 0.5. Sample size was determined to be 45 participants per group. To account for drop-out, 103 participants were recruited from the Trondheim community to participate in the study. All experimental procedures were approved by the Regional Committee for Medical Health and Research Ethics (2018/1310). The study was carried out following all relevant guidelines and regulations in Norway. Prior to the study, participants provided their written informed consent and were screened for contraindications. Participants were compensated for their time at a rate of 100 NOK per hour. Of the initial 103 participants, 5 were unable to complete the task because of motion sickness or fatigue. An additional 7 were excluded due to difficulty understanding the instructions and learning the task, and 1 participant was excluded due to participation in a related study. Therefore, 90 participants entered the analysis with 45 participants in each group (MG: 20 female, age range 20 - 34 years, mean age 25.2 years; GG: 23 female, age range 19 - 33 years, mean age 24.4 years). The MG was collected before the GG so that time point of sign-up for the study determined if a participant belonged to the MG or the GG. By collecting the MG prior to the GG we were able to match movement speed and exposure time between groups.

### Paradigm overview

The experimental paradigm consisted of a series of tasks on a desktop PC and in VR. Participants were first trained to use the motion platform and remote control (MG) or use the gamepad (GG). They were then familiarized with the task space through an interactive instructions task. Participants completed a pre-navigation forced choice test on a desktop PC. Afterwards, they entered a navigation training task and completed a series of angle and distance estimations. They concluded the experiment by completing the post-navigation forced choice task. Next we will explain the quantity space participants navigated. Then we will elaborate the experimental set-up as well as the tasks and analysis.

### Quantity Space

Participants were asked to navigate a two-dimensional abstract quantity space. This space was defined by the amount of circles and the amount of rectangles. Thus, a position in the space is defined as a point in a coordinate system defined by the number of circles and the number of rectangles (see figure 1). Each shape axis spanned from 0 - 200 shape quantities which translates directly into conceptual meters: the area of the quantity space was 200 x 200 conceptual meters. To match each position in visual complexity, we added triangles as filler shapes. This way each position had 400 items. The specific positions of the shapes was randomized in the visual field. Hence, a location appeared slightly different each time it was visited.

To indicate heading direction, we applied a unique color code to each shape dimension and the placeholder shapes. The color code ranged from red to blue, and all colors were matched in luminance. The color of the dimension 1 and dimension 2 shapes scaled with the cosine and sine of the heading angle respectively. It was balanced whether the circles or rectangles (and their corresponding sets of placeholder shapes) were assigned dimension 1 or dimension 2 (see figure 1 **b** for an illustration). Participants learned that if shapes were red (positive cosine or sine) and they moved forward, the quantity of this shape would increase. If the shapes were blue (negative cosine or sine), the quantity would decrease. If the shapes were purple (cosine or sine at zero), no change in the given shape quantity would occur. For an example of the color code, see the quantity space video S1 in the supplementary information (also available on the OSF, see Additional information).

### Experimental set-up

The virtual environment was programmed in Vizard (version 5.7, WorldViz LLC), a python-based VR software. To allow participants to physically navigate the conceptual space, we employed a novel VR set-up consisting of an omnidirectional motion platform (Cyberith Virtualizer) and a head mounted display (Oculus Rift CV1). The HMD had a resolution of 1080 x 1200 pixels and a refresh rate of 90 Hz. In the motion platform, participants were secured in a harness. The harness was attached to a rotatable ring, affording the ability to freely rotate. Participants wore low-friction overshoes which allowed them to slide their feet across the baseplate as they stepped forward. The virtualizer generated a speed value from the sliding motion across the baseplate. Orientation and speed were measured at 90 Hz.

In the MG, the motion platform was used to navigate the space. The speed and ring orientation readings controlled the participant’s position, and the HMD orientation readings were used to update the angular color code. In the GG, an Xbox One controller was used to navigate the space. The left stick on the controller determined the angular velocity while the right trigger determined the translational velocity. The translational velocity for both the GG and MG was smoothed by 4 frames (0.044 s).

### Tasks and Analysis

Participants completed three tasks including navigation training, distance and direction estimation as well as a two alternative forced choice task. All tasks are described in detail in the following sections. See figure 2 for an illustration. The instructions for all tasks avoided spatial language as much as possible. For example, participants were told to “create patterns” rather than to “navigate”. We instructed participants to change color and add shapes to create the pattern. All analyses were calculated in Matlab R2019a or RStudio 1.2.5.0.33 with R-3.6.3^54^. For all analysis the alpha-level was set to 0.05 unless otherwise stated. We further used Bayes factor analysis and will follow the convention for Bayes factor cut-offs and interpretation given in^55, 56^.

**Figure 2.**
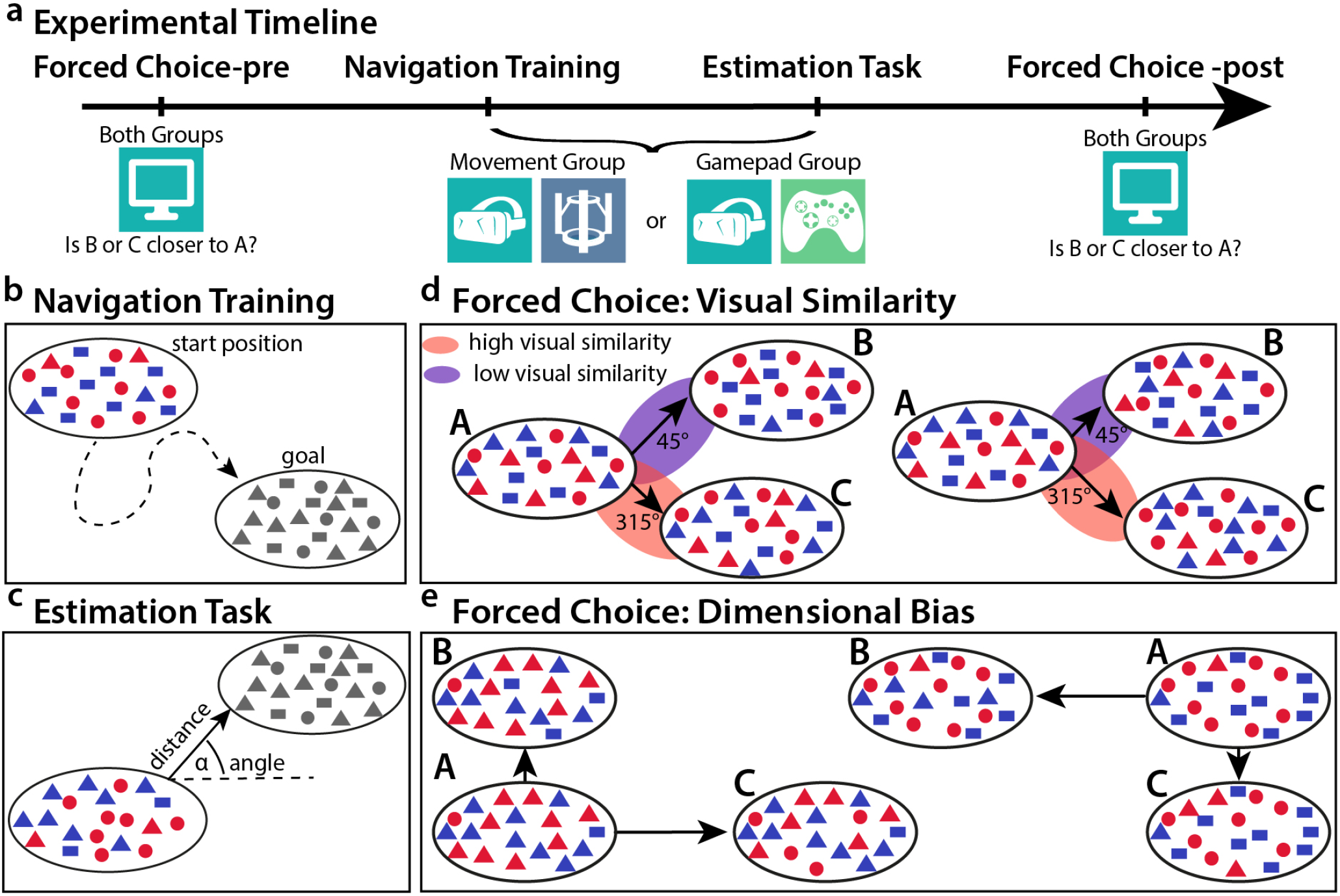
Tasks. **a** Experimental timeline. All participants conducted the experiment in the following order: forced choice pre, navigation training, distance and direction estimation task, forced choice post. The forced choice tasks were conducted on a computer screen while the other two tasks were group specific, either conducted on the motion platform or with the use of a gamepad. **b** In the navigation training, participants were shown a goal position and were then asked to navigate to it. When they thought they had reached the goal, they were instructed to press a button in order to receive feedback. **c** The estimation task required participants to estimate the angle and the distance between two given positions. They were first shown the goal position, then the start position. Then they were asked to turn in the direction of the goal. Finally, they used a slider to indicate the relative distance between the two positions. The slider ranged from the identical position to the furthest position possible. **d** Forced choice task to investigate visual similarity. Participants were shown three positions A, B and C at the same time. They had 5 seconds to decide if B or C was closer to A. We tested positions close to the center of the quantity space. B and C were positioned at different distances from A indicated by longer and shorter arrows, so that there always was a correct choice. The angles between positions A-B and A-C were chosen to maximize and minimize visual similarity according to the item overlap distribution described in figure 3. This allowed us to assess if participants were influenced in their judgments by visual similarity over distance. The left combination presents a trial in which high visual similarity (red) was aligned with the correct choice (shorter arrow), while the right combination presents a trial in which low visual similarity (purple) was aligned with the correct choice (shorter arrow). **e** Forced choice task for dimensional bias. Here, we chose positions that were close to the maximum (upper right corner) or the minimum (lower left corner) of item quantities, i.e. we showed positions with a lot of circles and rectangles, or positions with very few circles and rectangles. By manipulating only one dimension in B and the other dimension in C, we tried to find a dimensional bias, i.e. if participants found it easier to see the differences in the quantity of circles (here X-dimension) than that of rectangles (here Y-dimension) or vice-versa. Distances between the positions are indicated by the length of the arrows.

#### Navigation Training

Participants performed a navigation training task in which they navigated between a start and a goal position (Fig. 2b). They were cued with an image of the goal location at the beginning of each trial, and they were able to retrieve this image at any point during the task. The shapes defining the goal were presented in black, as to not prime the participants with a color code. Participants first completed a series of training trials, and later a set of testing trials. For the initial 5 training trials, participants were notified when they were in range of 10 percent of the goal location along both dimensions (100% being the total length of a dimension). For the subsequent 3 training trials, participants indicated when they reached the goal location with a button press. An error percentage was calculated by dividing the euclidean distance from the response location to the goal location by the maximum distance in the space. The error percentages were assigned into 5% increments from 0 to 100%, with each error range corresponding to a point score. Feedback was provided out of a maximum of 20 points, and displayed for 5 seconds. Along with the point value, a feedback message was shown to the participant depending on the error percentage.

For all training trials, participants were given unlimited use of a hint button. The hint indicated the distance to the goal in each dimension, and what color the participant should set the shapes. For the subsequent testing trials, the hint function was disabled. Participants had to complete a minimum of 14 testing trials. If this minimum number was met, participants in the MG continued until the time in the task reached 54 minutes. If the participants met the minimum number of testing trials in the GG, they continued until they completed 28 trials. This was done in order to keep the mean number of trials consistent between groups. All testing trials were self-paced and there was no time limit. Goal locations were distributed across the entire space and were at least 30 conceptual meters from the start position. The navigation was continuous, meaning participants used the indicated goal location of the previous trial as the new starting position.

Mean movement speed for the MG was 9.004, se = 0.458, for the GG’s mean movement speed was 9.099, se = 0.039. The speed was calculated using all testing trials of the task. Each trial was broken into segments of continuous movement. We excluded stationary periods and segments when the participant was viewing the goal or hint. If a distance of at least 10 conceptual meters was traveled within a segment, the mean velocity for the segment was calculated. This mean velocity was then calculated across all valid segments for each participant, and finally among participants. One subject in the GG never traveled a continuous 10 conceptual meters without pausing. This participant was therefore excluded from the calculation. We did not exclude this participant from the study, as the participant was not an outlier in other analyses. The movement speed corresponds to the number of items added/deleted per second during movement. Further, we added a slight fading function to the appearance of the individual shapes to ensure participants perceived the space as continuous. Additional tasks such as indicating responses and viewing messages were enabled by an Oculus Remote for the MG, and buttons on the Xbox One controller for the GG.

To assess performance in the navigation task, we looked at three main variables: accuracy, path length and navigation time. Accuracy was calculated by creating a random distribution of 1000 possible end-positions and calculating the proportion of positions that are further away from the true goal than the indicated end-position. This was done for each trial and each participant. An accuracy of 0.95 thus means that 95 percent of the random positions are further away and thus corresponds to very high accuracy. Excess path length was calculated for each trial by subtracting the optimal path length from the taken path length. This difference was normalized by dividing it by the optimal path length. Navigation time was the time participants spent navigating, excluding goal retrievals and breaks from the trial time. General performance was assessed using t-tests against chance for each group individually as well as non-paired two-sided t-tests to compare groups as implemented in R (function t.test). As supporting evidence for group comparisons, Bayes factor analyses were conducted to reveal more detailed information and hidden trends. Bayes factor analysis allows us to compare two models quantifying how much more evidence we have for one model over the other. Here we compare the null model (there is no difference between groups) with the alternative model (the groups differ). Thus we can also find evidence for equality of the groups, not just the absence of an effect. We used the BayesFactor package in R with the ttestbf function. We adhere to standard Bayes factor interpretation and cut-offs as given in^55, 56^.

We hypothesised that, similarly to observations of positional memory in physical space^41, 57^, accuracy in close proximity to boundaries would be higher than for positions in the center. For this, we correlated distance to the border with accuracy for each participant and then tested the correlation values on a group level against zero.

We assessed potential dimensional biases in goal perception by calculating the mean error in goal location for each participant for each of the two dimensions: circles (C) and rectangles (R), and compared them on a group level with a paired t-test. Additionally, we compared the two groups with a non-paired t-test on the difference between the circle and rectangle dimension.

We looked at learning during the navigation training by fitting general linear models (GLM) for each participant individually. We then extracted the t-statistic for the regressor of interest for each participant. Since the t-statistic does not follow a normal distribution, we used a sign-flipping permutation method to create a null-distribution. This entails randomly sign-flipping the t-statistics and calculating the mean of the new distribution. This procedure was repeated 10,000 times. The true mean of the t-statisic distribution was then compared to the sign-flipped mean-distribution. We could then calculate the proportion of means that are larger (or smaller) than our true mean, resulting in a p-value that we assessed using an α-level of 0.05. Z-values are created by subtracting the p-value from 1 and computing the inverse of the normal cumulative distribution function.

#### Distance and Direction Estimation Task

After participants had completed the navigation training, we tested whether they accurately represented distance and directional relations within the conceptual space (figure 2c). To this end, we asked participants to estimate the angle and distance between two cued locations. The goal location was presented in the same manner as in navigation training task, and could be viewed a maximum of 2 times with each view limited to 10 seconds. The first view occurred at the beginning of each trial, and the second view could be used by the participant at any point. After the first view of the goal, participants were given the starting location. They had to first estimate the direction from that location to the goal location. Participants indicated what color the shapes should be by turning their head in the correct direction (MG) or using the controls on the gamepad (GG). They indicated their response with a button press, and the color of the shapes was locked for the remainder of the trial. Participants then estimated how long it would take to create the target pattern from the start pattern by using a virtual slider. We used temporal distance instead of spatial distance since spatial language was avoided in the instruction of the task. The slider was set on a scale from 0 seconds to the time it took the participant to traverse the maximum distance in the space. The slider contained 29 increments from position 0 to position 28. For each trial, the initial slider position was always set to the center. Participants completed 3 practice trials using randomly generated locations. They then completed 36 testing trials. Trials evenly sampled locations separated by one of four distance bins (20-180 conceptual meters) and 9 direction bins (40 degree spacing). All test positions were at least 10 conceptual meters apart from each other as well as from the axis limits in order to span the entire space. Testing trials were organized into 4 blocks of 9 trials. No feedback was given in this task.

For the analysis, we correlated distance estimates with the true distances for each participant and tested the resulting correlation values on a group level against 0. We then compared the two groups against each other, first with a traditional t-test and second with Bayes factor analysis to reveal possible trends in the data. Since the abstract space allows us to look at distance and visual similarity between positions separately, we created linear regression models for each participant modelling the estimated distance with the true distance and the visual similarity. Visual similarity was expressed by the normalized number of shared shapes between two positions. We standardized visual similarity by calculating visual similarity between the center position and each position on a radius of 100 vm away from the center, then normalizing all values. A detailed explanation of visual similarity can be found in figure 3. Color did not factor into visual similarity here, since all tasks took measures to control for color in the comparisons, either by only comparing stimuli with the same color code or by presenting the stimuli in all black. We thus created a “look-up” table for visual similarity as a function of angle, independent of distance. With the same permutation method as described above, we compared the t-statistics for the two regressors.

**Figure 3.**
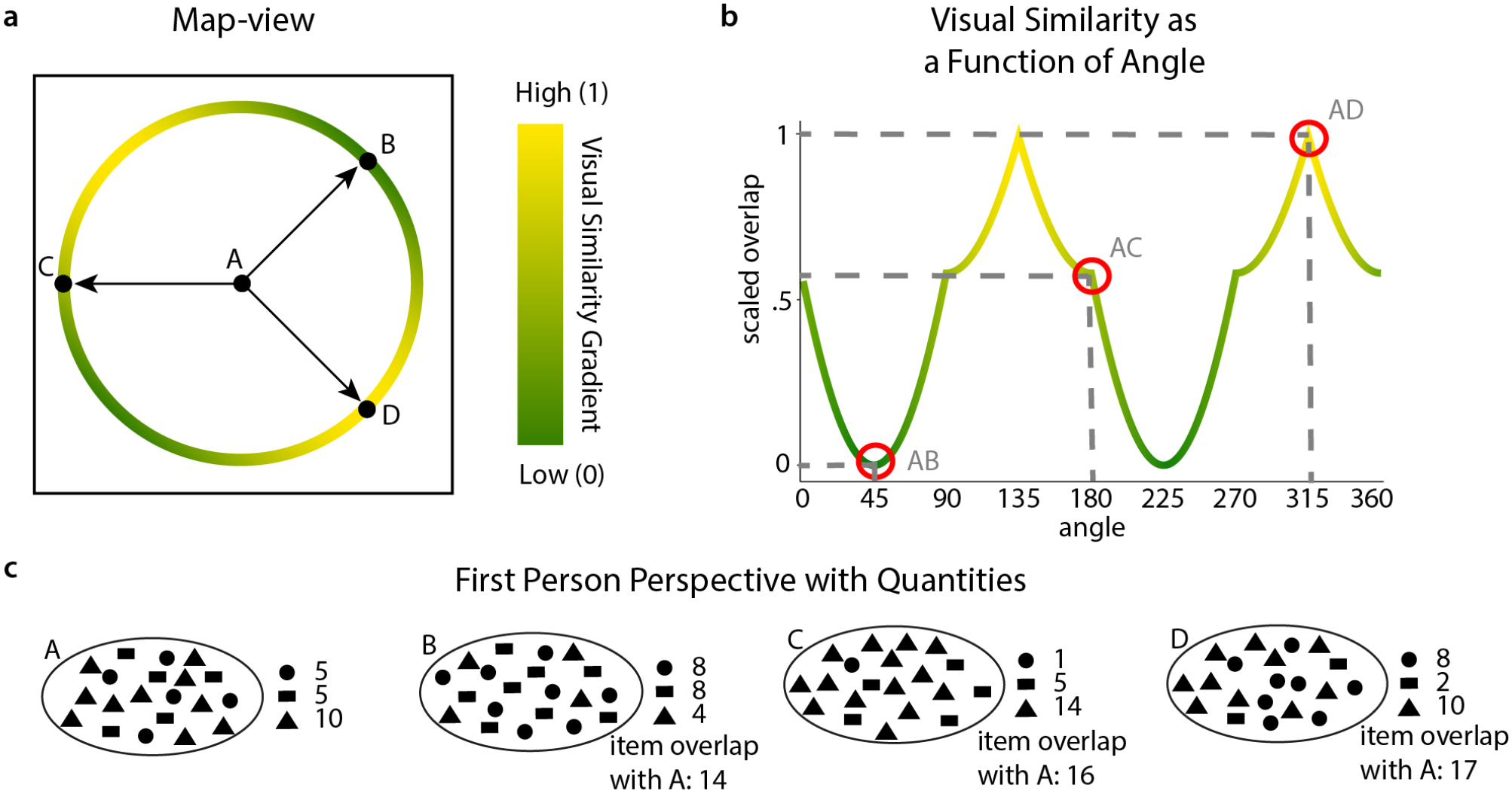
Visual Similarity. **a** Map-view. Representation of the space as given in figure 1. Shown are 4 positions, A, B, C and D. Positions B, C and D have the same distance to A. The circle on which they lie shows the angle dependent visual similarity distribution in a color gradient from yellow (high visual similarity) to green (low visual similarity). B has the lowest visual similarity to A, D has the highest. **b** Visual similarity as a function of angle. Visual similarity was calculated for all positions on a circle with radius 100 virtual meters and the center position (as in the example in **a** between A and all positions on the circle around it including B, C and D) by calculating the total amount of items shared between the positions. This distribution was then scaled to range between 0 (lowest visual similarity) and 1 and expressed as a function of angle. Items shared are independent of the color code that was used to encode direction (i.e. the red to blue scale). Highlighted are the visual similarities for the angles AB, AC and AD. **c** Down-sampled (5 %) representation of the first-person perspective when standing at the locations A, B, C and D as presented in **a**. For this illustration we excluded the color code, since all tasks that measure visual similarity controlled for color. Visual similarity was thus both independent of distance as well as of color and can be purely described as a function of angle (as in **b**). For each position we included the quantities of items. Here visual similarity as total item overlap is given in respect to A for positions B, C and D.

We analysed the angles by calculating the mean absolute angular error over all 36 trials for each participant followed by a t-test against chance (90 degrees) on a group level. We further compared groups with a t-test and Bayes Factor analysis to elaborate on the results. Next we wanted to establish if the distance and/or the visual similarity between positions had an effect on the angular error. Again we fitted a linear regression model for angular error for each participant, adding distance and visual similarity as regressors.

#### Forced Choice Task

We designed a 2-alternative forced choice task to test for effects of visual similarity and dimensional bias on distance perception. Participants completed this task both prior to the navigation training and at the conclusion of the experiment. With this method, we could determine whether the effects of visual similarity and dimensional bias changed as a result of navigation training. The forced choice test consisted of 216 testing trials with an additional 5 practice trials. Testing trials were organized into 4 blocks and each trial had a 5 second time limit. Between each testing trial, a message was flashed for 0.2 seconds indicating either that the response was recorded or that the participant did not answer in the allotted time. Participants did not receive feedback about their performance during testing trials. In the task, three locations were displayed: A, B, and C. Participants had to choose which of the two locations (B or C) was closer to the goal (A). The goal was presented at the top center of the screen, while the choice locations were presented in the lower left and right corners. Participants indicated their choice of B or C by pressing the left and right arrows on a keyboard. The color code for all three positions was the same in order to rule out any effects of color similarity. The specific settings for the forced choice task were determined during piloting and chosen in a way that participants performed slightly but significantly above chance. This way we could control if participants understood the instructions correctly. The forced choice test was divided into 2 sub-tasks. One sub-task tested for a dimensional bias among the shapes, while the other tested whether distance judgments scaled with the Euclidean distance or the visual similarity between the presented locations. In both tasks, we also tested which distance differences participants could accurately detect. The trials for both sub-tasks were intermixed and occurred in a random order.

To determine whether perception of closeness were affected by visual similarity beyond Euclidean distance between positions, we sampled choices that were at directions maximally similar (135 degrees and 315 degrees) and maximally dissimilar (45 degrees and 225 degrees) in terms of visual similarity (figure 2d). This is possible since the number of items shared between two positions is not only dependent on the distance but also on the angle between positions (see figure 3). The goal was placed in the center of the space 90 to 100 conceptual meters away from the axis limits in each dimension. One choice was distorted by 10 conceptual meters in one similarity condition (similar or dissimilar) while the other choice was distorted by 60, 90, or 120 conceptual meters in the alternative similarity condition. Locations were presented in a color code corresponding to 45 degrees or 225 degrees. In the visual similarity sub-task, we had 96 trials with 16 repetitions per each sampled distance condition.

Along with testing the effect of visual similarity, we tested for a bias of angular alignment and color priming. We aimed to determine whether the directional color code that participants learned could prime them to choose a certain location. To test this, we presented half of the trials in the dissimilar condition with a color code that was aligned with the correct answer’s true direction, and the other half in a color code that was misaligned. This was only possible in the dissimilar condition because we used color codes corresponding to 45 degrees and 225 degrees. Note that all three positions A, B, and C were always presented in the same color code, to make it easier to focus on the shapes only.

We also tested if there was a dimensional bias among the quality dimension shapes. To this end, we sampled positions in the lower left and upper right hand corner of the space where there was a dominance of placeholder and quality dimension shapes, respectively (figure 2e). This dominance stems from the logic of our quantity space: in the upper right corner, both quantity dimensions are near their upper limit, meaning participants saw a lot of circles and rectangles but only very few triangles. In the lower left corner, both quantity dimensions were near their lower limit, with only very few circles and rectangles but a lot of triangles present on the screen. Visually, differences in very low quantities are more salient. We therefore hypothesised that a dimensional bias would be accessible in these extreme cases. In this task, the goal was placed at a random location between 4 - 10 conceptual meters away from each axis limit for either corner. The choices were each distorted from the goal in a different dimension. One location was always distorted from A by 5 conceptual meters, while the other location was distorted from A by either 15, 30, or 55 conceptual meters in the alternate dimension. This created a layout in which one choice was always 10, 25, or 50 conceptual meters closer to A than the alternate choice. The dimensional bias sub-task consisted of 120 trials with 60 repetitions for both the upper and lower corner, and 10 repetitions per each of the 6 distance difference scenarios. The locations in this task were presented in a color code corresponding to 135 degrees and 315 degrees, with a random jitter of 10 degrees in either direction.

Analysis was run in R using the anova_test function from the rstatix toolbox. The data were collapsed across the different distances tested to maximize the number of trials. Further, performance or accuracy here is defined as percentage correct. To investigate the effect of visual similarity we used the data from the first sub-task and ran a three-way mixed ANOVA with one between factor of group (MG or GG) and two within factors for time point (pre and post) and visual-similarity (high and low). We checked ANOVA assumptions for outliers, normality and homogeneity, correcting for sphericity in our analysis. The exclusion of two potential outliers identified in these assumption checks did not change the interpretation of the statistical analysis and we thus report statistics based on data from all participants.

To investigate the effect of color priming, we looked only at trials in which the low-visual similarity option was the correct choice. We again used a three-way mixed ANOVA with one between factor (group) and two within factors for time point and alignment (aligned or misaligned with correct choice). Excluding outliers from the analysis did not affect the results.

For the second sub-task (the one investigating the possibility of a dimensional bias), we calculated a four-way mixed ANOVA with one between factor of group (MG or GG) and three within factors of time point (pre- or post-training), shape (circles or rectangles) and corner (lower left or upper right). Again, we identified extreme outliers that opposed ANOVA assumptions of normality. Since excluding the outliers did not change the results, we decided not to exclude them from this analysis.

## Results

### Navigation Training

#### General Performance

Here, we assessed the general performance of the participants in the navigation training to ensure participants in both groups understood the task and navigated in the instructed way. Firstly, we looked at accuracy, which is a measure of how close an indicated goal location is to the true goal location in comparison to a random distribution of possible locations. Both groups performed significantly above chance in indicating the goal location (chance-level = 0.5, mean_MG_ = 0.8623, t_MG_ = 33.6084, df_MG_ = 44, p < 0.001, mean_GG_ = 0.8641, t_GG_ = 38.6638, df_GG_ = 44, p < 0.001) but did not differ from one another (t = 0.1266, df = 86.4404, p = 0.8995). Bayes factor analysis revealed moderate evidence for the equality of the groups (BF_10_ = 0.2223). The mean trial duration for the MG was 116 seconds differing significantly from trial durations of the GG with a mean of 89 seconds (t = −3.3006, df = 81.792, p = 0.0014). The difference between groups is also strongly supported by Bayes factor analysis (BF_10_ = 22.5455). Excess path lengths did not differ significantly between groups (t = −1.9457, df = 82.107, mean_MG_ = 1.2594, mean_GG_ = 0.9517, p = 0.0551) with Bayes Factor analysis providing only anecdotal evidence against the null hypothesis (BF_10_ = 1.1476).

As has been found in physical space navigation^57, 58^, we expected to find an effect of distance to the boundary, i.e. accuracy should increase closer to the boundary. For both the MG and GG, we found significant negative correlations for distance to border and accuracy (mean_MG_ = −0.1929, t_MG_ = −5.646, df_MG_ = 44, p_MG_ = 1.115e^−06^, mean_GG_ = −0.2043, t_GG_ = −7.2935, df_GG_ = 44, p_GG_ = 4.245e^−09^) but no difference between groups (t = 0.2586, df = 84.747, p = 0.7965). Participants in both groups thus showed higher accuracy for goal positions at the borders than in the center, matching findings in physical space navigation.

Finally, we tested for a dimensional bias in the indicated goal location. A dimensional bias would mean participants perceive differences in quantities at different thresholds for the circle and the rectangle dimension. We found no significant dimensional bias in the MG (mean_C_ = 24.5346, std_C_ = 8.7333, mean_R_ = 23.4573, std_R_ = 8.6842, t = −1.1111, df = 44, p = 0.2726) but found a significant bias in the GG, with lower error in the rectangle dimension than in the circle dimension (mean_C_ = 25.6617, std_C_ = 10.5541, mean_R_ = 22.2035, std_R_ = 7.3661, t = −29786, df = 44, p = 0.0047). When comparing the two groups against each other with a non-paired t-test of the difference between error in the circle dimension and in the rectangle dimension we found no significant difference (t = −1.5741, df = 85.2889, p = 0.1192). The dimensional bias in the GG but not in the MG will be discussed below.

#### Learning

As a second step we included a separate analysis on the navigation training data, aiming to quantify participants’ learning behaviour. Specifically, we looked for improvements in navigation time, path lengths and accuracy. Navigation time and path lengths can be viewed as measures for efficiency and certainty, accuracy as a measure of precision.

To assess learning over the course of the training periods, we predicted navigation times from trial numbers in participant-specific linear regression models (figure 4**ab**). Trial number negatively predicted navigation time in the MG (p = 0.0001, z = 3.719) and the GG (p = 0.0001, z = 3.719). This indicates that participants became faster with more experience, thereby demonstrating learning. When comparing both groups using a non-paired, two-sided t-test we found no significant difference (t = −0.0377, df = 87.919, p = 0.97). Bayes Factor analysis provides moderate evidence for the equality of groups (BF_10_ = 0.2209). Taken together, these results suggest both groups show an equal amount of improvement in their navigation durations.

**Figure 4.**
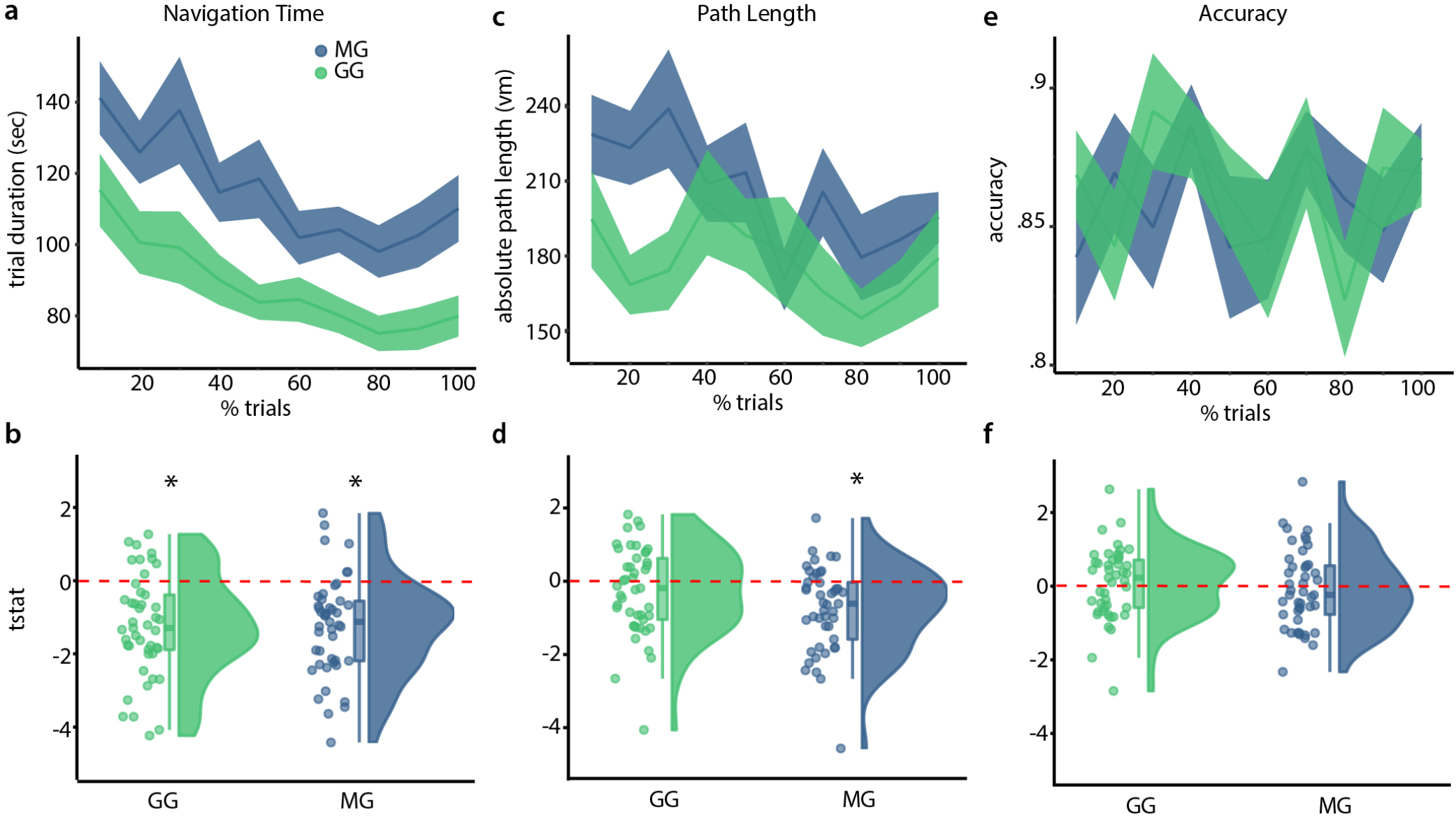
Navigation Training Results. **a**. Development of navigation time over the duration of the experiment. Since participants varied in the number of trials, we binned the trials in 10 bins for each participant and plotted the mean and standard error of trial duration for each bin across all participants. Note that we did not conduct any tests on the binned data, the plots are for visualization purposes only. **b**. t-statistics for each participant for predicting navigation time from trial number. Here, we use raincloud plots^59, 60^ that provide individual data points, box-plots with median marking as well as hinges indicating the 25^th^ and 75^th^ percentiles and a half-violin plot for a density distribution visualization. **c,d** Development of excess path length over the duration of the experiment equivalent to navigation time plots in **a** and **b**, respectively. **e,f** Development of accuracy over the duration of the experiment, equivalent to the plots in **a** and **b**, respectively. Green indicates participants from the gamepad group, blue from the movement group. Red dotted lines indicate chance level, * indicates a significant difference from the chance-level. See text for full statistics.

Next, we predicted path lengths from optimal path lengths and trial numbers(figure 4**cd**). We found that trial number negatively explained path lengths for the MG (p = 0.0001, z = 3.719) but not for the GG (p = 0.0674, z = 1.495), suggesting that the MG improved in finding the most direct path, while the GG did not improve. Testing for group differences in learning using a non-paired two-sided t-test, we failed to see a difference (t = −1.9764, df = 87.865, p = 0.0512). In addition, using Bayes factor analysis to elaborate on this trend we found only anecdotal evidence for a difference between groups (BF_10_ = 1.2086). As shown in the general performance section, mean excess path lengths did not differ between groups. This suggests that the absence of a learning effect in the GG was due to more accurate navigation in early trials, making it difficult to compare learning effects.

Lastly, we looked at accuracy scores, quantifying the precision with which participants navigated to target positions (see Methods, (figure 4**ef**)). Here, we modeled accuracy from trial numbers and observed no significant effect for either the MG (p = 0.6831, z = −0.4764) or the GG (p = 0.2926, z = 0.5458). We next compared learning between groups, first using a non-paired, two-sided t-test finding no significant difference (t = −0.7247, df = 87.314, p = 0.4706) and second using Bayes factor analysis (BF_10_ = 0.2782), revealing moderate evidence for the equality of groups. Accuracy thus remained stable over the experiment, which was likely due to participants being allowed to look at the goal as often as they liked, where participants would match their current position to the goal position.

### Distance and Direction Estimation task

In the estimation task, participants judged the angle and distance between pairs of positions (36 trials), which allowed us to probe their ability to plan goal-directed trajectories through the quantity space. The estimation task was conducted right after the navigation training (see figure 2**a**).

To test if participants could accurately estimate distances between a start and a goal location, we compared true distances with their distance estimates and found significant positive correlations in both the MG (t = 16.083, df = 44, mean Pearson r = 0.4968, p < 0.001) and the GG (t = 19.288, df = 44, mean Pearson r = 0.5353, p = 0.001). We then compared the two groups against each other, finding no significant effect (t = 0.9269, df = 87.01, p = 0.3565). These results provide evidence, that both groups performed the distance estimation task successfully but did not differ from one another. Bayes Factor analysis supports the absence of a difference between groups (BF_10_ = 0.3221). Next, we wanted to assess factors that could drive distance estimates. We used visual similarity and true distance to predict distance estimates and found that both distance (p_MG_ = 0.0001, z_MG_ = 3.719, p_GG_ = 0.0001, z_GG_ = 3.719) and visual similarity (p_MG_ = 0.0001, z_MG_ = 3.719, p_GG_ = 0.0001, z_GG_ = 3.719) were significantly contributing to participants’ estimates (figure 5). This illustrates that both true distances as well as visual similarity affect judgement in this task.

**Figure 5.**
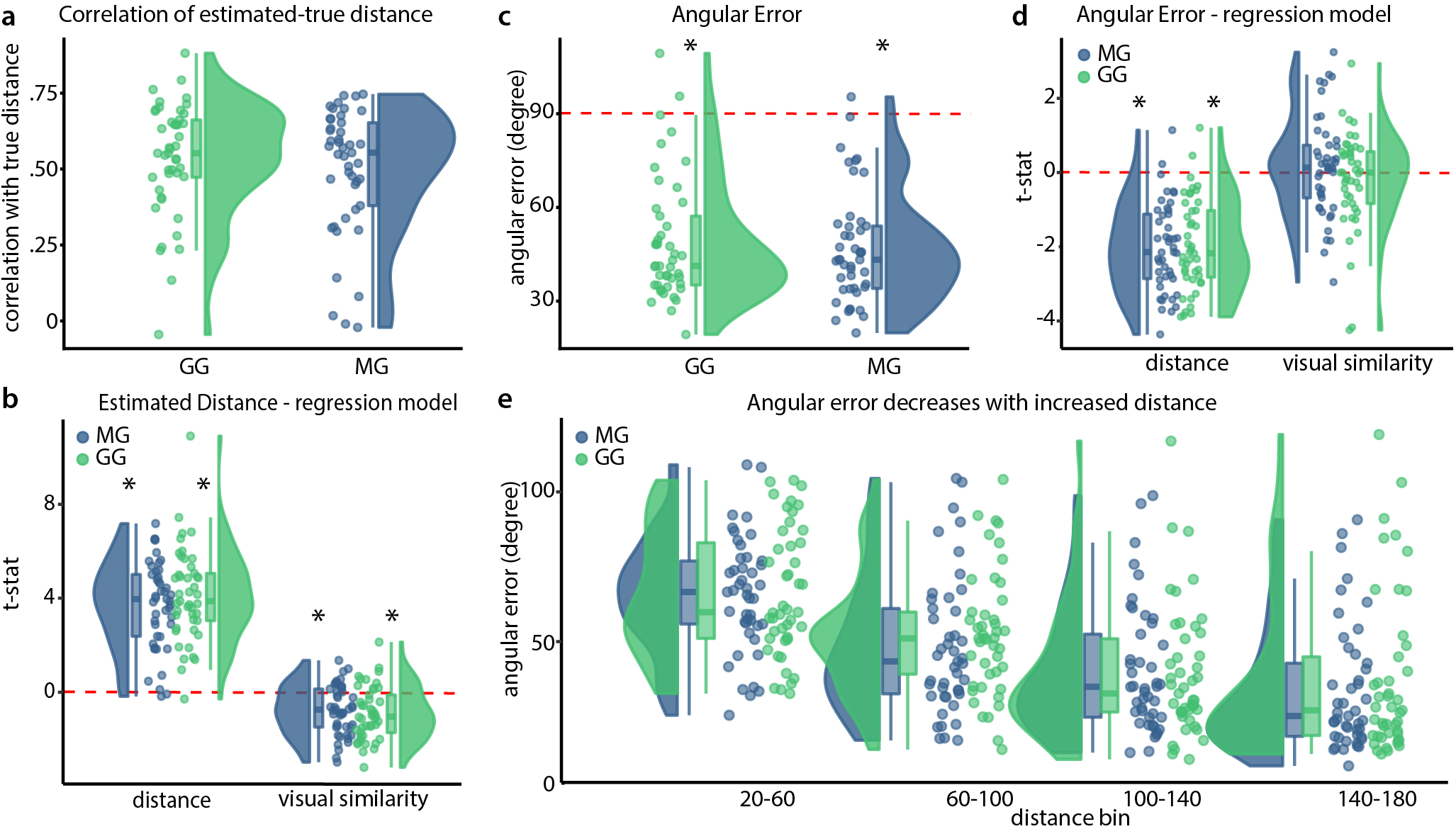
Distance and Direction Estimation task results. **a** Correlation of true and estimated distances. We find significant positive correlations for both groups but no difference between groups. **b** T-statistics from regression model for distance estimation. Regression models are fitted individually for participants, t-statistics are then compared in a permutation test. See text for an exact explanation of the procedure. Both visual similarity and distance influence distance estimation in both groups. **c** Absolute Angular Error. We find low errors for both groups, significantly below chance error level. **d** t-statistics from regression model for angular error for distance and visual similarity. Angular errors are smaller with increased distance, but show no linear relation to visual similarity. **e** Illustration of relationship between angular error and distance between positions. The goal and start location had 4 different distances as indicated by the distance bins on the x-axis. Performance in the angle estimation increased with increased distances. Green indicates participants from the gamepad group, blue from the movement group. We use raincloud plots to visualize individual data points, box plots with marks for median as well as 25^th^ and 75^th^ percentile percentiles and a half-violin plot for a density distribution. Red dotted lines indicate chance level. See text for exact statistics.

In addition to estimating the distance of a given start position to a goal location, we asked participants to estimate the direction to the goal location. By regarding both components we could assess the accuracy of the mental representation of the abstract space. Both groups showed absolute angular errors smaller than expected from chance level behaviour (t_MG_ = 17.599, df_MG_ = 44, mean_MG_ = 46.9936, p_MG_ < 0.001, t_GG_ = 16.64, df_GG_ = 44, mean_GG_ = 48.9885, p_GG_ < 0.001), showing they could estimate directions with good accuracy (figure 5). We further compared groups, but found no significant difference (t = 0.5019, df = 87.175, p = 0.617). Using Bayes Factor analysis to elaborate on these results (BF_01_= 0.2466), we found slightly lower angular errors for the MG than for the GG (see^55^ for conventions concerning Bayes Factor interpretation). Next we tried to predict absolute angular error from distance and visual similarity and found distance to be a significant predictor (p_MG_ = 0.0001, z_MG_ = 3.719, p_GG_ = 0.0001, z_GG_ = 3.719), while visual similarity was not a significant predictor (p_MG_ = 0.7814, z_MG_ = −0.7769, p_GG_ = 0.1637, z_GG_ = 0.9794). See figue 5 for illustrations of these results. Direction unlike distance perception was thus independent from visual similarity, but absolute angular error did scale with distance; suggesting an integration of the two features.

### Forced Choice Task

The forced choice task was performed before as well as after the navigation training and estimation task. We therefore have two time points: “pre” and “post”. We further designed trials in two different ways, to test for effects of visual similarity and color priming (sub-task one) and dimensional bias and symmetry of space (sub-task two, see methods).

#### Effect of visual similarity

In this part of the forced choice paradigm, we chose positions B and C to be either aligned or misaligned with the visual similarity distribution, to test if participants showed a bias towards choosing the visually similar choice even if this was not the choice closer to the probe. Further, we aimed to investigate if a possible bias was reduced with training, i.e. participants learned to base their choice more on distance than on visual similarity. We performed a three-way mixed ANOVA with one between factor of group (MG or GG) and two within factors for time point (Pre or Post) and visual similarity (high or low). We found a main effect of time point (F = 19.974, df = 88, p = 2.33e^−05^) and of similarity (F = 356.811, df = 88, p = 1.03e^−32^) providing evidence that performance increased after training and that high-visual similarity affects distance judgements. We observed no effect of group (F = 0.015, df = 88, p = 0.902) and no interactions. See figure 6 for an illustration and supplementary table S1 for the full ANOVA table. Participants thus showed a visual similarity bias, which was not reduced after training.

**Figure 6.**
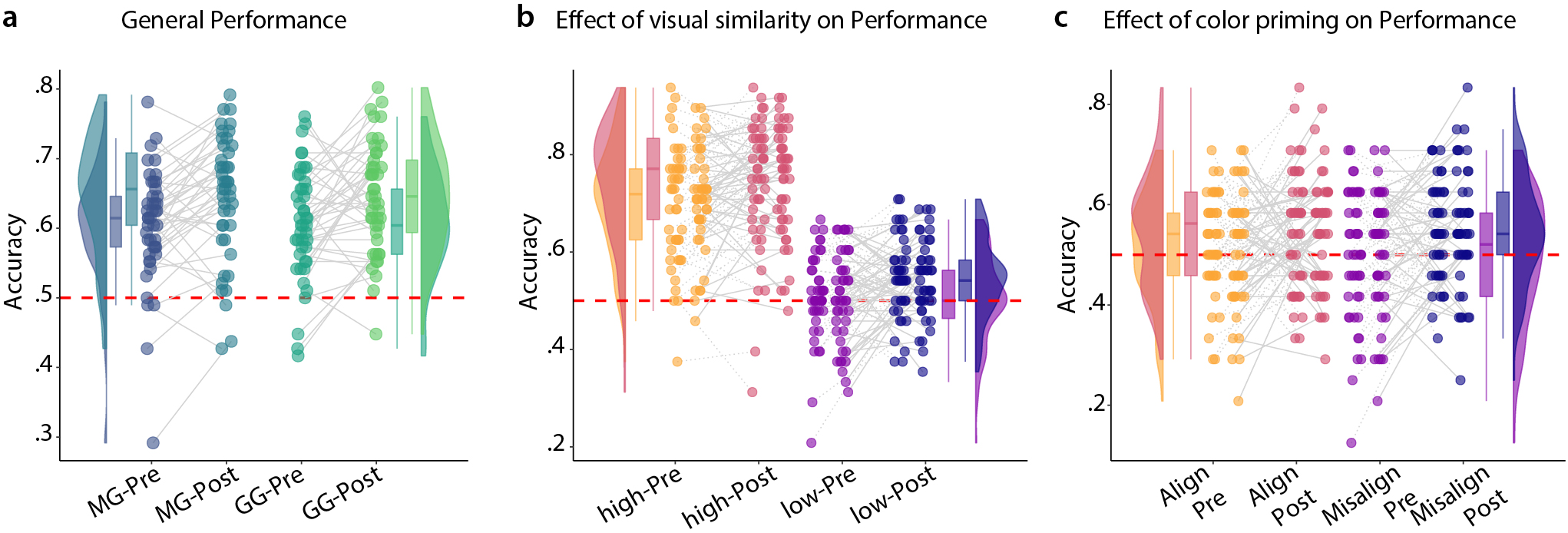
Results from the forced choice task investigating visual similarity and color priming. **a** General performance split up by groups and time point: pre or post. Mixed ANOVA revealed a main effect of time-point but not of group. **b** Main effect of visual similarity on performance. Violin and bar plots contain data from both groups. Scatter plots in each condition are split into two columns: the first column for the movement group, the second for the gamepad group. There was no main effect of group. Lines connect individual measures. **c** Illustration of the absence of an effect of color priming in the data. As in **b**, the two columns of identically colored scatter dots illustrate participants’ performance in the movement group and the gamepad group respectively. Note that only trials in the low-visual similarity condition were used in this analysis, therefore fewer trials factored into the analysis.

Next we used only half of the trials (those in the low-visual similarity condition) to assess an effect for color priming or angle priming. We hypothesised that if the color applied (color encodes direction) is aligned with the angle between AB or AC, the aligned position may be a more likely choice. We again used a three-way mixed ANOVA with one between factor (group) and two within factors for time point and alignment (aligned or misaligned with correct choice). We found no main effect of alignment (F = 0.125, df = 88, p = 0.725), group (F = 0.067, df = 88, p = 0.796) nor any interaction effects (supplementary table S2). We again found the main effect of time point (F = 10.1369, df = 88, p = 0.002). We could therefore not find any evidence for color priming induced by learning.

#### Effect of dimensional bias

The second sub-task of the forced choice task compared biases participants may have for the two dimensions defined by the rectangles and circles. A dimensional bias would entail, that differences in quantities may be more accurately perceived in either the circle or the rectangle dimension. Therefore, we chose position B to differ from A only in one dimension, and position C to differ from A in the other dimension. Further, we tested positions in the upper right and lower left corner of the space to determine if participants’ performance changed when they had to rely on the filler items instead of the circle and rectangle dimensions and if there was a systematic difference in performance in these corners of the space. Positions in the upper right corner were defined by very high amounts of circles and rectangles (low amounts of fillers). Positions in the lower left were defined by very low amounts of circles and rectangles (high amounts of fillers). We calculated a four-way mixed ANOVA with one between factor of group (MG or GG) and three within factors of time point (pre or post), shape (circles or rectangles) and corner (lower left or upper right). We found a main effect of time point (F = 59.989, df = 88, p = 1.53e^−11^), which suggest learning, and a main effect of shape (F = 7.69, df = 88, p = 7.00e^−03^) which suggests a dimensional bias. We found no other main or interaction effects (see supplementary table S3). In post-hoc analyses, we computed pair-wise comparisons between all subgroups looking for an effect of shape to understand the dimensional bias. We found a significant effect of shape only in the upper-right corner in both pre (df = 44, p = 0.017) and post (df = 44, p = 0.034) for the GG; however, this does not survive correction for multiple post-hoc tests (supplementary table S4). Since we find the subtle dimensional bias in pre in the GG but not in the MG, our experimental manipulation cannot explain the dimensional bias. See figure 7 for an illustration.

**Figure 7.**
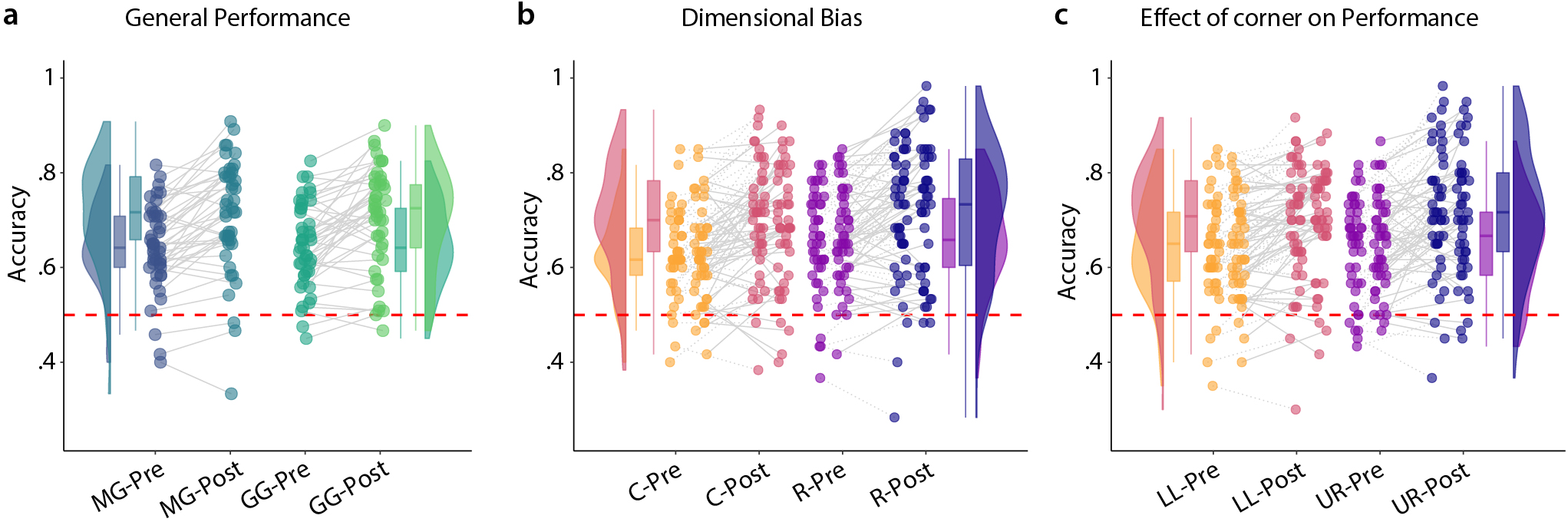
Results from the forced choice task investigating dimensional bias and corner. **a** General performance split into the movement group and the gamepad group as well as pre and post. We found a main effect of time point, but no group effect. **b** Illustration of dimensional bias analysis. Box plots and violin plots contain the data from both groups, scatter points are split into two columns: on the left the movement group on the right the gamepad group. We found a slight advantage for rectangles (R) over circles (C) but no interaction effects and no main effect of group. **c** Illustration of performance in the lower left (LL) and upper right (UR) corner of the space in pre and post. Corners of the space refer to the positions in the quantity space in a Cartesian coordinate system: lower left as low values both along the X- and Y-dimension (few circles and rectangles) while upper right refers to high values in the X- and Y-dimension (many circles and rectangles). Again, density and bar plots contain data from both groups while scatter points are organised in columns (the movement group on the left, the gamepad group on the right). We found a main effect of time point but no other effects.

## Discussion

In this study, we created a first-person view abstract space navigation task. We closely matched aspects of physical space, such as the number of dimensions, mode of navigation and informational content. Our aim was to establish a new paradigm that combines both theories of cognitive spaces and embodied learning: an abstract space that uses first-person navigation, contains both positional as well as directional information, can be learned by physical exploration and can easily be extended. We used the paradigm to investigate a key question of the field: If we build up cognitive maps of any type of knowledge, and if physical interaction enhances learning, can we navigate an abstract space in the same manner we navigate the real world? Furthermore, does physical movement improve learning?

Our results suggest that participants can learn the representation of an abstract environment both through physical navigation and through the use of a gamepad. We find trial durations shortening over time in the navigation task as well as accurate performance in angle and distance judgements. Participants demonstrate learning in the forced choice task with increased performance after navigation training. We did not, however, find strong differences between groups, suggesting that our embodied learning manipulation did not aid in the facilitation of an accurate representation of the space.

Along with observing learning in the three sub-tasks, we found certain biases that resemble navigational biases during spatial exploration of the physical world. For example, in the navigation training we found that the estimation of position was more accurate near the borders than in the center of the space. This effect becomes also evident when comparing the distance judgements in the forced choice test: during piloting we found that for center positions we needed larger differences of distances (50, 80 and 110) than for positions in the corners (10, 25, and 50). Yet, performance in the corner positions was still higher than in the center positions (see supplementary information text as well as tables S5 and S6). This is consistent with findings from studies testing positional memory in navigation paradigms^57, 58^.

Additionally, in the angle estimation task, visual similarity did not explain angular error, but the larger the distance between two positions the smaller the angular error. Participants thus made smaller errors in the angle estimation when the distances between positions were large. This is not surprising since the general direction from one position to the other, i.e. the differences in the quantities are more apparent when positions are far apart. For example, a participant is given a starting position A and has to estimate the angle to position B. When B is far away, the participant is able to reduce the possible angles considerably by judging if more or fewer circles as well as more or fewer rectangles are needed. Thus the 360 degree decision space is reduced to a 90 degree decision space. If position B on the other hand is close to A and potentially within the perceptual threshold, the decision space cannot be reduced as easily and the estimation is more difficult^61^. In typical spatial navigation research, this can be easily explained when researchers consider the visual angle also known as the angular size of a goal location: an object that is far away from you has a smaller visual angle on your retina than an object that is close to you. In our task, the same rules apply, when we take into account that participants do not have a dot-like representation of the individual positions but rather a field-like representation. This is evident in the perceptual thresholds of a position in the space (see forced choice paradigm). Estimating the exact angle to the goal thus becomes more difficult if the goal is close to you, compared to when it is far away.

In this study, we show that participants can navigate an abstract space in the same manner they navigate physical space, even showing similar behavioural biases. But, what are the implications of such a paradigm? For one, navigation is a well-trained, intuitive behaviour. Here we show that it can be applied to the acquisition of knowledge beyond the layout of a physical space. This opens up a new avenue of learning techniques: providing abstract information in a spatial manner, that in turn can be physically navigated. By mapping relationships between abstract stimuli onto a spatial representational format, learners could acquire the knowledge in an intuitive manner. Imagine for example, learning about politics or deciding whom to vote for. Politicians (or parties) can be represented as individual points (or areas) in the large n-dimensional space of opinion. We could for example, represent politicians along three dimensions: the degree to which they like to invest in science, the degree to which they see global warming as a problem and the degree to which they support capitalism. Using a spatial layout makes it very easy to estimate which two politicians have the most similar opinions (are the closest in space) and which differ the most. You could also navigate to your own position in this high dimensional space and then see which politicians are close to you, by potentially viewing the dimensions that are most important to you or using dimensional reduction techniques. Large amounts of information, i.e. all the different positions of the different politicians would therefore become spatially intuitive and easily comparable.

Spatial learning strategies are already established in the field of memorization: the method of loci uses an imagined mind palace, in which the to be remembered information is placed in different rooms. When the knowledge needs to be recollected, participants imagine navigating through their mind palace visiting the positions at which they previously placed the items. This memorization strategy has shown to be very effective^62^ not only in imagination^63^ but also in combination with a virtual environment^64^.

Language also uses several spatial metaphors to describe abstract relationships. “You couldn’t be further from the truth”, “A comes before B”, “It is close to midnight", are examples of spatial language that describe non-spatial content. These types of metaphors have been shown to shape the way we think^65^ pointing at the inherent link of how we learn and how knowledge is represented in our brain^66^.

We constantly learn new, complicated concepts and making use of spatial representations with advanced VR techniques provides an enticing opportunity. Beyond the possibilities for learning, our paradigm could be useful for investigations of knowledge representations: do we represent knowledge in a spatial manner and do we use mechanisms from spatial navigation to access and manipulate non-spatial information? One interesting part of spatial navigation is the flexibility of switching between allocentric and egocentric reference frames. Relations or distances between landmarks and other positions in the space are preserved in both allocentric and egocentric maps^66^. In our study, we find evidence for such a preservation of relations between positions well as the integration of directional information and distances. This suggests that participants do acquire a cognitive map of the quantity space, even when they only use a first-person perspective to navigate the space. The paradigm itself is easily extendable with another dimension or additional “rooms” (by adding a new kind of shape, once one dimension is at its maximum). We could also include landmarks, obstacles, or distortions to test the limits of abstract cognitive maps.

Why was there no clear difference between groups? We had hypothesised that participants in MG would show clear differences in performance to GG, since they used physical movement to navigate and thus had multiple senses providing information that could be integrated and potentially reduce error.

One possibility is that our task provided too much information: path integration and multi-sensory integration was not necessary, since all information of position and direction was constantly available in the current view. Visual accessibility has been pointed out as a factor that renders physical movement for accurate formation of a cognitive map^30^ unnecessary. Our space was therefore easily solvable. Still we found perceptual (visual) biases in performance that could have been reduced after embodied learning. Here it is possible that effects of our group manipulation are only detectable with more training distributed over multiple days.

Further differences in performance may only be evident on a longer time-scale: Johnson-Glenberg et al. (2016)^67^, trained students in physics concepts by varying the degree of embodied learning in the knowledge acquisition. They found that performance in a physics test right after training did not differ between groups but when they tested the students again one week later they found a significant effect of embodiedness: the higher the level of embodiment at encoding the better the retention of the knowledge. For future studies that use embodied learning approaches we would therefore recommend post-tests to assess retention of knowledge gained.

In our task, we are careful not to rule out an effect of embodied learning due to some limitations our set-up. The motion platform recordings could be noisy, which resulted in less smooth navigation. Furthermore, participants were more exhausted using the motion platform. The movement on the motion platform deviated from a natural walking motion in that the participants had to slide their feet across the sensors and lean into the harness. This resulted in slower and more effortful navigation. These effects may account for the finding that the MG exhibited longer trial durations despite having similar path lengths. We also found a dimensional bias in the GG but not in the MG, both in the navigation training as well as in the forced choice task. This indicates that the GG demonstrates a consistent dimensional bias that is not diminished by careful consideration as is done in the navigation training. In the forced choice task, participants only had 5 seconds to decide, while in the navigation training participants were free to spend any amount of time as well as to compare the goal and current location as often as they liked. Careful consideration of the quantities could therefore overrule a purely perceptual dimensional bias that we assessed in the forced choice paradigm. Since this bias was already present in the pre-training forced choice task, at which point the group manipulation had not yet begun, we can unfortunately not explain the bias, since it seems to be a coincidental effect, not based on our experimental manipulation. However, when comparing the two groups directly, we find no significant differences. We therefore reason that any other results of group differences should not be dependent on a dimensional bias present in one group.

Our finding that an abstract space can be navigated both with and in the absence of physical movement introduces future questions. One potential direction is to investigate how training over a longer duration could elicit an effect of embodied learning. Furthermore, it could be of interest to adapt this into an imaging study in an effort to understand the neuronal correlates of our behavioural measures. The use of a map-like representation of knowledge could also be interesting for research in generalization and transfer of knowledge^41, 42^. For example, do we transfer relationships between items made in one space into another? Both the method of loci, as well as spatial language of metaphors point to space as a useful tool to understanding and remembering complicated relations^40, 65^. We suggest that intuitive spatial behaviour paired with constantly improving VR technology could be a useful tool for learning. Future research should also aim to test if participants inherently organize abstract knowledge in a spatial manner, or if applying a spatial format to abstract knowledge through experimental manipulation is necessary for such an organization.

In conclusion, our paradigm shows that first-person navigation of abstract spaces is possible, using not only button presses but also physical movement. Participants are able to flexibly use color and quantities as directional and positional information. Such a paradigm can be used to assess the usefulness of multi-sensory integration in the form of embodied learning (potentially on a longer time-scale)^8^, landmark- and boundary-based navigation in abstract spaces^46^ (i.e. prototyping), short-cuts, inferences and transferability of knowledge between different cognitive spaces. Physical exploration of an environment is an intuitive and already well-trained task. Presenting knowledge in a map-like fashion that can be explored thus poses an interesting opportunity for teaching as a new “learning strategy": walking and creating connections between objects in an abstract space.

## Acknowledgements

This work is supported by the European Research Council (ERC-CoG GEOCOG 724836). C.D.’s research is further supported by the Max Planck Society, the Kavli Foundation, the Centre of Excellence scheme of the Research Council of Norway – Centre for Neural Computation (223262/F50), The Egil and Pauline Braathen and Fred Kavli Centre for Cortical Microcircuits, and the National Infrastructure scheme of the Research Council of Norway – NORBRAIN (197467/F50).

## Author contributions statement

D.K., J.B. and C.D. originally designed the study and its hypothesis. N.S. programmed the task and collected the data, under the supervision of D.K.. D.K. analysed the data and wrote the first draft of the manuscript. R.K., J.B. and C.D. supervised the work. All authors contributed to the final version of the manuscript.

## Additional information

### Accession codes

Data and code is available at: https://osf.io/nvu4t/

