## Supplementary Information for "An immersive first-person navigation task for abstract knowledge acquisition"

#### This PDF file includes:

Supplementary text
Tables S1 to S6
Legend for Movie S1
SI References

#### Other supplementary materials for this manuscript include the following:

Movie S1

### Supporting Information Text

**Additional settings for the Navigation Training.** To ensure that we did not introduce systematic differences between groups by keeping our trial structure flexible in the navigation training, we used Bayes factor analysis to provide evidence for the equality of groups (1). We calculated the mean distance between start and goal over all navigation training trials for each participant and compared the groups using the BayesFactor package (version 0.9.12-4.2) in R. Bayes factors compare the likelihoods of two models against one another. Here, we tested the alternative hypothesis (means differ between groups) against the null model (there is no mean difference) and found moderate evidence ($bf_{10} = 0.232913$) for the null hypothesis (2). This indicates that we did not introduce systematic differences between groups in the design of our navigation training trials.

**Comparison of performance in the two sub-tasks of the forced choice task.** The forced choice task was split into two sub-tasks: a task evaluating the effect of visual similarity and the other evaluating the effect of dimensional bias. In the visual similarity sub-task, positions in the center of the space were tested, while in the dimensional bias sub-task, positions near the upper right and lower left corner of the space were compared. During piloting, we evaluated which distances should be compared to reach performances above chance level. This led to larger distances in the center of the space than in the corners. For the forced choice performance, we use a three-way mixed ANOVA with 1 between (group) and 2 within-subject factors (time point, i.e. pre or post and taks (i.e. visual similarity or dimensional bias). We find a main effect of time point ($F = 60.264$, $p = 1.41e^{-11}$) and of task ($F = 44.145$, $p = 2.45e^{-09}$) as well as an interaction effect between task and time point ($F = 4.983$, $p = 2.80e^{-02}$). We find no other main effects or interaction effects (see table S5). In table S6 we display mean and standard errors for the different combinations to illustrate that even though participants were tested on smaller distances in the dimensional bias task they reached higher accuracy levels than in the visual similarity task.

**Table S1. Three-way mixed ANOVA with one between factor for group, and two within factors for time point and visual similarity.**

| | Effect | DFn | DFd | F | p | p<.05 | ges |
|---|---|---|---|---|---|---|---|
| 1 | group | 1 | 88 | 0.01 | 9.02e-01 | | 8.18e-05 |
| 2 | visual similarity | 1 | 88 | 356.81 | 1.03e-32 | * | 4.79e-01 |
| 3 | time point | 1 | 88 | 19.97 | 2.33e-05 | * | 3.40e-02 |
| 4 | group : visual similarity | 1 | 88 | 0.26 | 6.10e-01 | | 6.75e-04 |
| 5 | group : time point | 1 | 88 | 0.03 | 8.73e-01 | | 4.60e-05 |
| 6 | visual similarity : time point | 1 | 88 | 0.00 | 9.55e-01 | | 5.11e-06 |
| 7 | group : visual similarity : time point | 1 | 88 | 0.10 | 7.57e-01 | | 1.55e-04 |

**Table S2. Three-way mixed ANOVA with one between factor for group, and two within factors for time point and alignment.**

| | Effect | DFn | DFd | F | p | p<.05 | ges |
|---|---|---|---|---|---|---|---|
| 1 | group | 1 | 88 | 0.07 | 7.96e-01 | | 2.40e-04 |
| 2 | alignment | 1 | 88 | 0.12 | 7.25e-01 | | 3.09e-04 |
| 3 | time point | 1 | 88 | 10.37 | 2.00e-03 | * | 3.00e-02 |
| 4 | group : alignment | 1 | 88 | 0.80 | 3.74e-01 | | 2.00e-03 |
| 5 | group : time point | 1 | 88 | 0.01 | 9.24e-01 | | 2.67e-05 |
| 6 | alignment : time point | 1 | 88 | 0.68 | 4.11e-01 | | 2.00e-03 |
| 7 | group : alignment : time point | 1 | 88 | 0.13 | 7.19e-01 | | 3.09e-04 |

**Table S3. Four-way mixed ANOVA with one between factor for group and three within factors for time point, shape and corner.**

| | Effect | DFn | DFd | F | p | p<.05 | ges |
|---|---|---|---|---|---|---|---|
| 1 | group | 1 | 88 | 0.00 | 9.85e-01 | | 1.92e-06 |
| 2 | shape | 1 | 88 | 7.69 | 7.00e-03 | * | 1.00e-02 |
| 3 | corner | 1 | 88 | 1.43 | 2.34e-01 | | 1.00e-03 |
| 4 | time point | 1 | 88 | 59.99 | 1.53e-11 | * | 5.10e-02 |
| 5 | group : shape | 1 | 88 | 1.90 | 1.72e-01 | | 2.00e-03 |
| 6 | group : corner | 1 | 88 | 0.29 | 5.94e-01 | | 2.99e-04 |
| 7 | group : time point | 1 | 88 | 0.26 | 6.12e-01 | | 2.32e-04 |
| 8 | shape : corner | 1 | 88 | 0.18 | 6.75e-01 | | 1.55e-04 |
| 9 | shape : time point | 1 | 88 | 0.94 | 3.35e-01 | | 4.60e-04 |
| 10 | corner : time point | 1 | 88 | 1.08 | 3.02e-01 | | 5.53e-04 |
| 11 | group : shape : corner | 1 | 88 | 0.29 | 5.93e-01 | | 2.53e-04 |
| 12 | group : shape : time point | 1 | 88 | 1.13 | 2.91e-01 | | 5.53e-04 |
| 13 | group : corner : time point | 1 | 88 | 0.78 | 3.78e-01 | | 4.02e-04 |
| 14 | shape : corner : time point | 1 | 88 | 0.89 | 3.48e-01 | | 5.53e-04 |
| 15 | group : shape : corner : time point | 1 | 88 | 0.06 | 8.03e-01 | | 3.88e-05 |

**Table S4. Post-hoc pairwise comparisons between all subgroups of the dimensional bias data from the forced choice task.**

| | time point | group | corner | shape1 | shape2 | n1 | n2 | statistic | df | p | p.adj | p.adj.signif |
|---|---|---|---|---|---|---|---|---|---|---|---|---|
| 1 | Post | not-walking | L | C | R | 45 | 45 | -1.18 | 44 | 0.24 | 0.24 | ns |
| 2 | Pre | not-walking | L | C | R | 45 | 45 | -1.61 | 44 | 0.11 | 0.11 | ns |
| 3 | Post | not-walking | U | C | R | 45 | 45 | -2.19 | 44 | 0.03 | 0.03 | * |
| 4 | Pre | not-walking | U | C | R | 45 | 45 | -2.49 | 44 | 0.02 | 0.02 | * |
| 5 | Post | walking | L | C | R | 45 | 45 | -0.75 | 44 | 0.46 | 0.46 | ns |
| 6 | Pre | walking | L | C | R | 45 | 45 | -1.62 | 44 | 0.11 | 0.11 | ns |
| 7 | Post | walking | U | C | R | 45 | 45 | 1.06 | 44 | 0.30 | 0.30 | ns |
| 8 | Pre | walking | U | C | R | 45 | 45 | -1.34 | 44 | 0.19 | 0.19 | ns |

**Table S5. Three-way mixed ANOVA with one between factor for group, and two within factors for time point and task.**

| | Effect | DFn | DFd | F | p | p<.05 | ges |
|---|---|---|---|---|---|---|---|
| 1 | group | 1 | 88 | 0.00 | 9.63e-01 | | 1.62e-05 |
| 2 | task | 1 | 88 | 44.15 | 2.45e-09 | * | 7.10e-02 |
| 3 | time point | 1 | 88 | 60.26 | 1.41e-11 | * | 7.00e-02 |
| 4 | group:task | 1 | 88 | 0.02 | 8.85e-01 | | 3.66e-05 |
| 5 | group:time point | 1 | 88 | 0.18 | 6.75e-01 | | 2.21e-04 |
| 6 | task:time point | 1 | 88 | 4.98 | 2.80e-02 | * | 4.00e-03 |
| 7 | group:task:time point | 1 | 88 | 0.07 | 7.98e-01 | | 5.34e-05 |

**Table S6. Mean performance as well as standard deviation and standard error split by group, task and time point.**

| | time point | group | task | mean performance | N | sd | se |
|---|---|---|---|---|---|---|---|
| 1 | Post | GG | visual similarity (center) | 0.64 | 45 | 0.08 | 0.01 |
| 2 | Post | GG | dimensional bias (corner) | 0.70 | 45 | 0.11 | 0.02 |
| 3 | Post | MG | visual similarity (center) | 0.64 | 45 | 0.09 | 0.01 |
| 4 | Post | MG | dimensional bias (corner) | 0.71 | 45 | 0.12 | 0.02 |
| 5 | Pre | GG | visual similarity (center) | 0.61 | 45 | 0.08 | 0.01 |
| 6 | Pre | GG | dimensional bias (corner) | 0.65 | 45 | 0.09 | 0.01 |
| 7 | Pre | MG | visual similarity (center) | 0.60 | 45 | 0.08 | 0.01 |
| 8 | Pre | MG | dimensional bias (corner) | 0.64 | 45 | 0.09 | 0.01 |

**Movie S1. Video illustrating the logic of the quantity space**

The video introduces the basic concepts of the abstract space in a stepwise manner. Captions on top of the video explain each step. Only when all three shapes (circles, rectangles and triangles) are presented, the space is complete. There are four different views, highlighting different aspects of the abstract space. **Allocentric Map**: A bird's-eye view of the abstract space. The x-axis represents circles and the y-axis represents rectangles. The white cursor indicates the participant's position within the space and the red line indicates the trajectory taken. **Heading Direction**: The direction the participant was facing and the corresponding color of the circles and rectangles. **Counter**: The amount of circles and rectangles displayed in the scene at each instance. This corresponds to the participant's coordinate position within the space. **Participant's View**: A depiction of the participant's view in the head-mounted display. The shapes appeared at random positions and were presented in an oval layout. Note that the three plots on the left were never shown to participants. **Captions: A**. When a participant walks forward in the space, shapes are added to the scene. **B**. The color of the shapes represents the participant's heading direction. When a participant turns, the color of the shapes changes along a gradient from red to blue. **C**. When the shapes are red, the participant is heading in a positive direction along the shape axis. The quantity will increase for the given shape unless the participant has reached the boundary of the space. **D**. When the shapes are blue, the participant is heading in a negative direction along the shape axis. The quantity will decrease for the given shape unless the participant has reached the boundary of the space. **E**. So far, the circle dimension has been at 0. If the participant moves towards the center of the space, they are moving along the circle dimension. When the color of a shape is purple, the quantity of the purple shape does not change. This is because the participant is not travelling along this shape axis. In this example, the quantity of rectangles does not change. However, the quantity of circles increases because the participant is heading in a positive direction along the circle axis. **F**. The circles and rectangles serve as the dimensions of the space. To ensure all positions had the same overall number of shapes, triangles were added as placeholders. Therefore, every position has 400 shapes. Each dimension has a corresponding set of triangles, and the color of the triangles matches the color of their respective dimension. This is now the full view of the quantity space. **G**. The heading angle determines which shapes are added or removed. When the participant is heading in a positive direction along the circle and rectangle axis, these shapes increase while the triangles decrease in quantity. When the participant is heading in a positive direction along the circle axis and a negative direction along the rectangle axis, the circles increase while their triangles decrease in quantity. The rectangles decrease while their triangles increase in quantity. Watching the counter, you will see that the total number of shapes remains constant at 400. **H**. The rate of change in shape quantity depends on the heading angle. Notice how first the circles change at a greater rate than the rectangles. This is because the participant is travelling a greater distance along the circle axis than the rectangle axis at this angle. When the participant turns, the rectangles change at a greater rate than the circles. This is because the participant is travelling a greater distance along the rectangle axis than the circle axis.
